# Generating Property-Matched Decoy Molecules Using Deep Learning

**DOI:** 10.1101/2020.08.26.268193

**Authors:** Fergus Imrie, Anthony R. Bradley, Charlotte M. Deane

## Abstract

An essential step in the development of virtual screening methods is the use of established sets of actives and decoys for benchmarking and training. However, the decoy molecules in commonly used sets are biased meaning that methods often exploit these biases to separate actives and decoys, rather than learning how to perform molecular recognition. This fundamental issue prevents generalisation and hinders virtual screening method development. We have developed a deep learning method (DeepCoy) that generates decoys to a user’s preferred specification in order to remove such biases or construct sets with a defined bias. We validated DeepCoy using two established benchmarks, DUD-E and DEKOIS 2.0. For all DUD-E targets and 80 of the 81 DEKOIS 2.0 targets, our generated decoy molecules more closely matched the active molecules’ physicochemical properties while introducing no discernible additional risk of false negatives. The DeepCoy decoys improved the Deviation from Optimal Embedding (DOE) score by an average of 81% and 66%, respectively, decreasing from 0.163 to 0.032 for DUD-E and from 0.109 to 0.038 for DEKOIS 2.0. Further, the generated decoys are harder to distinguish than the original decoy molecules via docking with Autodock Vina, with virtual screening performance falling from an AUC ROC of 0.71 to 0.63. The code is available at https://github.com/oxpig/DeepCoy. Generated molecules can be downloaded from http://opig.stats.ox.ac.uk/resources.

## 1. Introduction

Virtual screening is a computational approach that is often used in early stage drug discovery to help find molecules that interact with protein targets with high affinity and specificity. Numerous prospective applications of virtual screening have been reported, reducing the cost and improving the hit-rate of experimental verification (e.g. 15).

There are a variety of datasets available for benchmarking virtual screening methods through retrospective validation. These sets consist of a collection of active and inactive molecules for a range of protein targets. Frequently used examples for structure-based virtual screening (SBVS) are DUD (6) and DUD-E (16), DEKOIS (1, 26), and MUV (19).

While experimentally-verified inactives represent the gold standard for dataset construction (11), suitable inactive molecules are not typically available. As such, using presumed inactives, known as decoys, is almost always necessary (20). There are efforts to construct sets using only known inactives (e.g. 19, 24); however, these are relatively small in size and are not yet suitable for training modern machine learning methods.

Using decoys introduces three main sources of bias: artificial enrichment, analogue bias, and false negative bias (20). Artificial enrichment captures the performance that can be attributed to the differences in chemical space between the active and decoy molecules. Analogue bias arises from limited diversity of the active molecules, while false negative bias describes the risk of active compounds being present in the decoy set, which could lead to an underestimation of the screening performance. To limit these biases, decoys should resemble active molecules as closely as possible while simultaneously minimising the chance of the compound being a binder. This is routinely achieved through chemical property matching and structure mismatching, with all frequently used datasets selecting decoys from a database of molecules (1, 16), typically ZINC (23).

However, property matching arbitrary actives is challenging and, despite improvements, still leads to substantial differences in molecular properties between actives and decoys (2). While some properties that should be matched can be relevant for binding, such as molecular weight, it should not be possible to discriminate actives from inactives from these properties alone. However, this is routinely possible on several widely-used datasets (22, 27). Hence close matching is essential for dataset construction to reduce over-optimistic retrospective testing that cannot be replicated prospectively.

In recent years, many machine learning methods have been trained and evaluated on these datasets (e.g. 7, 29). The reported results show that these methods substantially outperform other methodologies such as empirical and knowledge-based scoring functions at SBVS.

Concerningly, some reports have suggested that a key driver of the performance of machine learning-based systems is hidden biases in the training data, such as physicochemical differences, and that these methods are not learning to perform molecular recognition (3, 22). Better decoy molecules are essential to remove the biases in datasets that are hindering the development of virtual screening methods.

The challenges of decoy design are in part due to the inherent limitations of matching to an explicit, fixed database of potential decoys. While virtual libraries such as ZINC (23) have grown considerably, they still represent only a tiny fraction of potential drug-like chemical space (17) and are insufficient for closely matching core chemical properties of many active molecules.

Wallach and Lilien (28) pioneered the use of a generative approach to construct virtual decoy sets for the original DUD (6) targets with tighter property matching than the decoys selected from ZINC. They used a rules-based algorithm employing a library of chemical building blocks and bridges to iteratively generate possible decoys. However, their method ignored synthetic feasibility and, despite clear improvements in property matching, has not been widely adopted.

Machine learning models for molecule generation have been proposed as an alternative to human-led design and rules-based transformations and have shown great promise in several molecular design tasks (e.g 21,31).

In this work, we describe DeepCoy, a deep learning method using graph neural networks, to generate decoy molecules. DeepCoy takes as input an active molecule and generates property-matched decoy molecules. This eliminates the need to use a database to search for molecules and allows decoys to be generated for the requirements of a particular active molecule and the user’s specification.

The properties can be chosen by the user depending on their objective, and in this paper we demonstrate the ability of DeepCoy to learn to produce decoy molecules with different sets of matched properties, highlighting the flexibility of our approach. We validated our generative model using two established SBVS benchmarks, DUD-E and DEKOIS 2.0. For all 101 DUD-E targets and 80 of the 81 DEKOIS 2.0 targets, our generated decoy molecules more closely matched the physicochemical properties deemed by the respective datasets to be non-informative for binding, improving property matching measured by DOE score by 81% and 66% for DUD-E and DEKOIS 2.0, respectively.

Finally, we demonstrate that the generated decoys are harder to distinguish from active molecules than the original decoy molecules with docking using Autodock Vina (25). This ability to substantially reduce bias will benefit the development and improve generalisation of structure-based virtual screening methods.

## 2. Methods

This work describes a novel approach using deep learning to propose molecules that match a set of features provided by the user. We achieve this with a generative model using graph neural networks. Our model makes no underlying assumptions regarding the nature of the properties that are to be matched, and relies only on a training set of paired molecules exhibiting the desired similarities.

### 2.1. Generative model

In order to generate decoys we use an adapted version of Imrie *et al.* (8), which was designed for linker generation. Imrie *et al.* (8) builds on the generative process introduced by Liu *et al.* (14) that constructs molecules “bond-by-bond” in a breadth-first manner. The most substantial differences with Imrie *et al.* (8) are the input data and goal of the generative process.

DeepCoy takes an active molecule as input and generates a new molecule that has similar physicochemcial properties but is structurally dissimilar. This is achieved by building new molecules in an iterative manner “bond-by-bond” from a pool of atoms. In this framework, the user is able to control the maximum number of heavy atoms in the molecules and, if desired, specific heavy atoms or partial substructures.

Minimal chemical knowledge is directly incorporated in our model; this takes the form of a set of permitted atom types and basic atomic valency rules which ensure the chemical validity of generated molecules. The model is required to learn all other decisions required to generate molecules.

Our method learns through a supervised training procedure using pairs of molecules (Figure 1). Inspired by Jin *et al.* (9), we frame decoy generation as a multimodal graph-to-graph translation problem. We train DeepCoy to convert graphs of active molecules into property-matched decoys under an augmented variational autoencoder setting, employing standard gated-graph neural networks (13) in both the encoder and decoder. DeepCoy implicitly learns which properties to keep constant and is not explicitly told which properties to match, nor their values. This provides a highly flexible framework, and makes it possible to learn from pairs of molecules without quantifying their similarity.

**Fig. 1.**
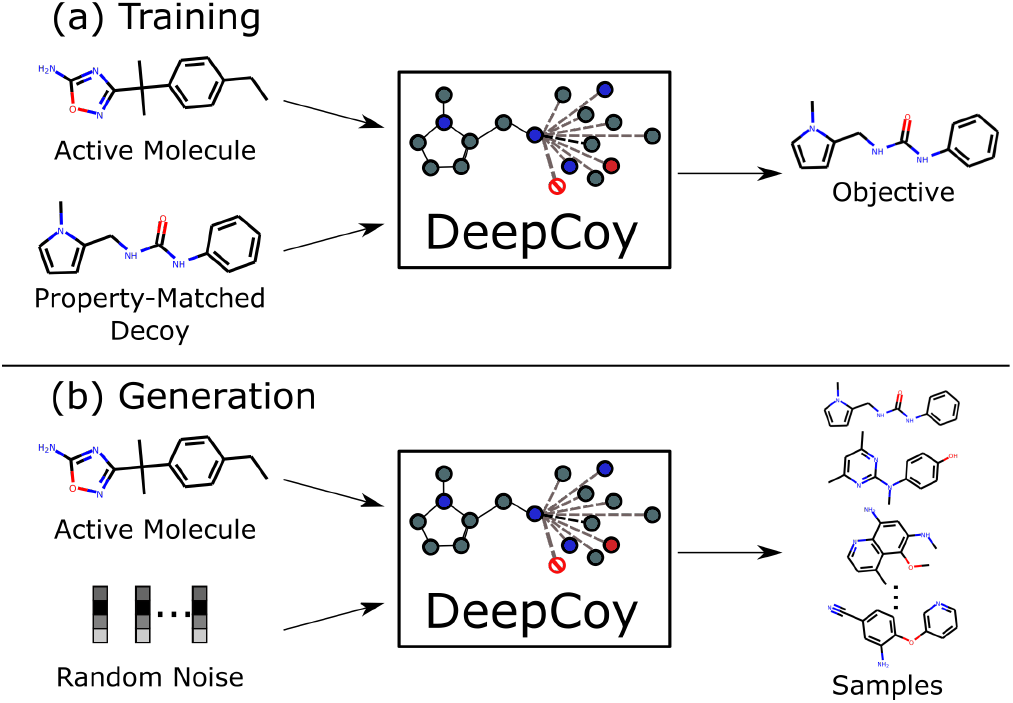
Illustration of training and generation procedures. (a) Pairs of structurally dissimilar molecules with similar physicochemical properties are provided as input. The model is trained to convert one molecule into the other from a combination of the encodings of both molecules. (b) At generation time, the model is given only the active molecule and is able to sample a diverse range of property-matched decoy molecules by combining the encoding of the active molecule with random noise.

We employed a training objective similar to the standard VAE loss, including a reconstruction loss and a Kullback-Leibler (KL) regularisation term:

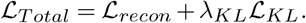

The reconstruction loss, 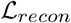, is composed of two terms resulting from the error in predicting the atom types and in reconstructing the sequence of steps required to produce the target molecule.

To improve the quality of generated molecules, we adopted a novel loss function that deviates from a standard cross entropy loss for the sequence of actions adopted by Imrie *et al.* (8) and Liu *et al.* (14). Instead of each step in the generative processes having equal importance, we reweighted the probabilities of actions by the frequencies of the induced subgraphs across the training set of molecules, leading to the revised cross-entropy loss:

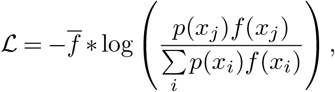

where *p*(*x_i_*) is the probability of choosing action *x_i_*, *f*(*x_i_*) is the reciprocal frequency of the induced local subgraph by taking action *x_i_*, the sum is over all permitted actions, and 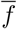 is the average of *f* over all permitted actions. This has the effect of reducing the chance of introducing local subgraphs that are not present in the training set. We observe that this change does not meaningfully affect the novelty of generated molecules compared to the standard cross-entropy loss.

For a more detailed description of the model, see Imrie *et al.* (8) and the Supplementary Information.

### 2.2. Training set

We constructed pairs of molecules to train our model from the 250 000 molecule subset of ZINC (23) selected at random by Gomez-Bombarelli *et al.* (5) as follows.

We first characterised compounds by their physicochemical properties. The properties can be selected by the user and we demonstrate the effectiveness of our framework using multiple sets of properties (described in Section 2.4). Pairs of molecules were constructed to satisfy the following criteria: (1) identical heavy atom count and counts of specific heavy atoms (C, N, O, S, Cl, F), (2) high similarity in property-space, and (3) low structural similarity. We measured similarity in property-space using the Euclidean distance between normalised property values and structural similarity by the Tanimoto similarity between the Morgan fingerprints (radius 2, 1024 bits, 18).

In order to create training sets for our large scale bench-marking experiments (see Section 2.4), we set the maximum permitted structural similarity between a pair of molecules at 0.15 and the maximum distance in property space to 0.20 for the assessment on DUD-E and 0.07 for DEKOIS 2.0. The thresholds were set to ensure roughly equal training set sizes and was as a result of the differences in properties to unbias. This resulted in a training set of 131,199 pairs for DUD-E and 103,170 for DEKOIS 2.0. We selected 1000 pairs for model validation, and used the remainder to train our model.

### 2.3. Assessment

Several metrics have been proposed to assess artificial enrichment and the risk of false negatives introduced by using putative decoy molecules. Vogel *et al.* (26) proposed the deviation from optimal embedding score (DOE score) and the doppelganger score to assess the quality of physicochemical matching of decoys and risk of introducing latent active molecules, respectively. These metrics are our primary way of assessing the generated decoy molecules.

The DOE score measures the quality of the embedding of actives and decoys in chemical space by employing a series of receiver operating characteristic curves (ROC curves) for each active calculated using the physicochemical properties of interest. The DOE score is the average absolute difference between these ROC curves and a random distribution. An optimal embedding of actives and decoys achieves a DOE score of zero, while complete separation in physicochemical space results in an DOE score of 0.5.

The doppelganger score captures the structural similarity between actives and their most structurally related decoys. We generated functional fingerprints (similar to FCFP6) using RDKit (12) for all compounds and evaluated the structural similarity between actives and decoys using the Tanimoto coefficient. For each decoy molecule, its doppelganger score is the maximum similarity across all actives. For each target, we report the mean doppelganger score over all decoys and the maximum structural similarity between an active and a decoy.

An alternate way to quantify the physicochemical property matching is via predictive models trained on such properties (22, 27). We trained 1-nearest neighbour (1NN) and random forest (RF) models on all possible subsets of the physicochemcial properties deemed non-informative for binding. We adopted 10-fold cross-validation on a per target basis and assessed performance via the area under the ROC curve (AUC ROC), following Sieg *et al*. (22). Other measures exist to assess bias in molecular datasets, such as AVE (27). However, rather than using a proxy, we chose to measure machine learning performance directly.

We also considered the virtual screening performance of docking using AutoDock Vina (25), specifically the smina (10) implementation. Ligands were docked against the reference receptor within a box centered around the reference ligand with 8 Å of padding. We used smina’s default arguments for exhaustiveness and sampling. We focussed our analysis on performance as measured by AUC ROC.

### 2.4. Large scale benchmarking experiments

We assessed our method using two of the most popular SBVS datasets, DUD-E (16) and DEKOIS 2.0 (1).

We trained a separate model for each of the datasets to demonstrate the flexibility of our method to learn to match different sets of properties. For DEKOIS 2.0, we used the same eight properties employed to construct the dataset (1). To assess whether our framework extends to a higher dimensional property space, we trained our model to match twentyseven properties for our assessment on DUD-E, instead of only the original six properties selected by Mysinger et *al.* (16). Training set construction is described in Section 2.2 and a complete list of physicochemical properties is provided in the Supplementary Information. Despite training for this broader array of properties and selecting the final decoys for the DUD-E set based on all 27 properties, we report results calculated using the original six DUD-E properties, unless otherwise stated. Not selecting the DeepCoy set optimally with respect to the original six DUD-E properties will result in inferior performance of DeepCoy, but will allow us to evaluate how our method performs when required to unbias a larger number of properties.

We filtered the active molecules in both datasets to exclude those containing rare atom types outside of the scope of our model (c. 1% of actives, see the Supplementary Information for a list of permitted atom types). This led to no actives for DUD-E target FPPS, so we excluded this target. For each active, we generated 1000 candidate decoys using DeepCoy. We then selected final decoy sets using a similar pipeline to DEKOIS 2.0. Generated molecules were initially filtered by the difference in heavy atom counts and maximum doppelganger score using an iterative procedure until at least 100 candidate decoys remained. The final decoys were selected from these candidate decoys in a greedy manner based on the sum of the normalised property difference and LADS score (1). We then compared the generated decoy sets to the original decoy sets using the metrics described in Section 2.3.

## 3. Results and Discussion

We assessed our ability to generate property-matched decoy molecules with varying requirements through two widely-used SBVS datasets, DUD-E and DEKOIS 2.0. For both sets, we generated new decoy molecules and compared these to the original set, assessing the generated molecules with respect to the same physicochemical properties used to select the original decoys. We show that:

- DeepCoy generated decoys substantially improve property matching compared to the original database decoys.
- DeepCoy generated decoys do not introduce additional risk of false negatives.
- DeepCoy generated decoys are harder to distinguish from active molecules than the original DUD-E decoys with docking using AutoDock Vina, despite being as structurally dissimilar from the active molecules as the original decoys.

Our results demonstrate that our framework is an alternative to database approaches for selecting property-matched decoy molecules, while offering full flexibility to the user regarding choice of specific properties and how to choose the final decoys from the generated molecules.

### 3.1. Physicochemical property matching

Across both DUD-E and DEKOIS 2.0, our generated decoy molecules more closely matched the physicochemical properties deemed by the respective datasets to be non-informative for binding than the original decoys (see the Supplementary Information for a full list of properties).

When selecting decoys based on the same properties as the original datasets, our generated decoys improved the DOE score by an average of 81% and 66%, respectively, decreasing from 0.163 to 0.032 for DUD-E and 0.109 to 0.038 for DEKOIS 2.0. In this setting, the DOE score was improved by using DeepCoy generated decoys for all 101 DUD-E targets (Figure 2) and 80 of the 81 DEKOIS 2.0 targets (Figure S1). The only DEKOIS 2.0 target that did not show an improvement in DOE score had DOE scores below 0.04, corresponding to an almost perfect embedding for both the DeepCoy and original decoy molecules. Finally, DeepCoy generated decoys achieved a DOE score below 0.1, indicating a close to optimal embedding (1), for 100 of the 101 DUD-E and 79 of the 81 DEKOIS 2.0 targets, while the original decoys only met this threshold for 32 DUD-E and 48 DEKOIS 2.0 targets.

**Fig. 2.**
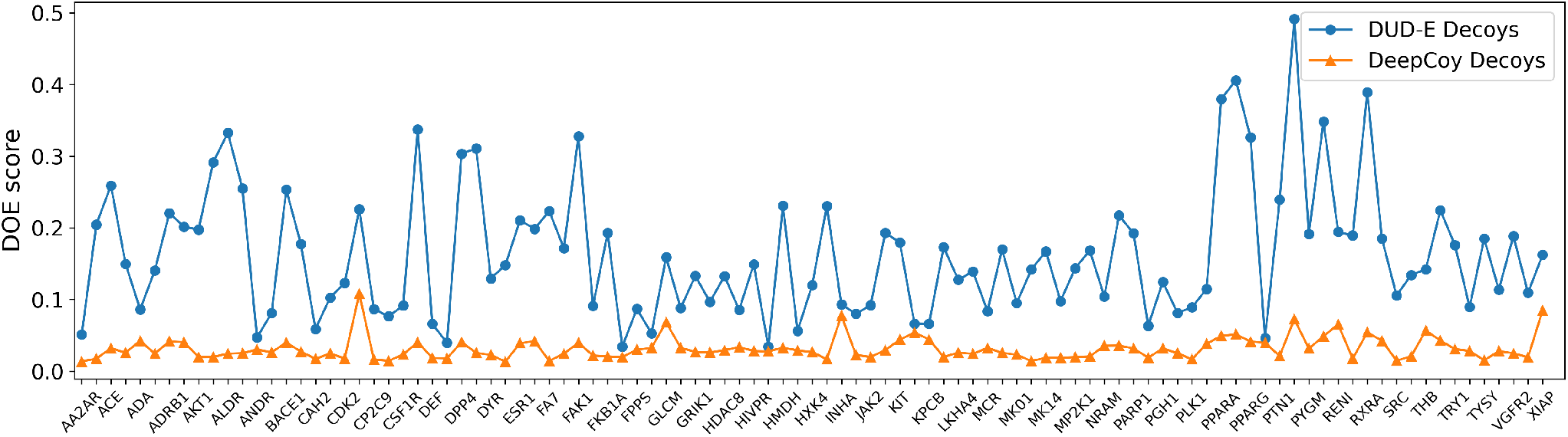
DOE scores of the original DUD-E set (blue) compared to the DeepCoy generated decoys (orange). For all targets, the DeepCoy generated decoys have lower DOE score (lower is better), with the average DOE score decreasing by 81% from 0.163 to 0.032. The x-axis displays each DUD-E target in the same order as they appear in the DUD-E database (http://dude.docking.org/targets), excluding FPPS. The targets with even indices are not labeled on the x-axis due to space limitations.

We selected our final decoy set for DUD-E using all 27 properties, rather than just the six used to construct the original dataset. The average DOE score of this set was 0.044, a comparable improvement of 73%, outperforming the original decoys for 97 of the 101 targets (Figure S2). Importantly, the DeepCoy decoys experienced no drop in performance when all 27 properties were included in the calculation of DOE score, with an average score of 0.040 (Figure S3). In contrast, the original decoys experienced a substantial decline to 0.220, proving matching this larger set is non-trivial. This demonstrates the ability of DeepCoy to scale successfully to a high-dimensional property space to unbias.

A similar improvement can be seen when assessing property matching via the ability of machine learning models to predict whether a compound is an active or a decoy when trained on the physicochemcial properties deemed non-informative for binding (Figures 3, S4). On the DUD-E set, using all 6 features, the median (average) AUC ROC decreased from 0.66 (0.66) to 0.57 (0.58) and 0.81 (0.80) to 0.70 (0.71) for the 1-nearest neighbour and random forest models respectively for the DeepCoy decoys compared to the original set. A similar reduction was observed when using any combination of the physicochemical properties (Figure 3).

**Fig. 3.**
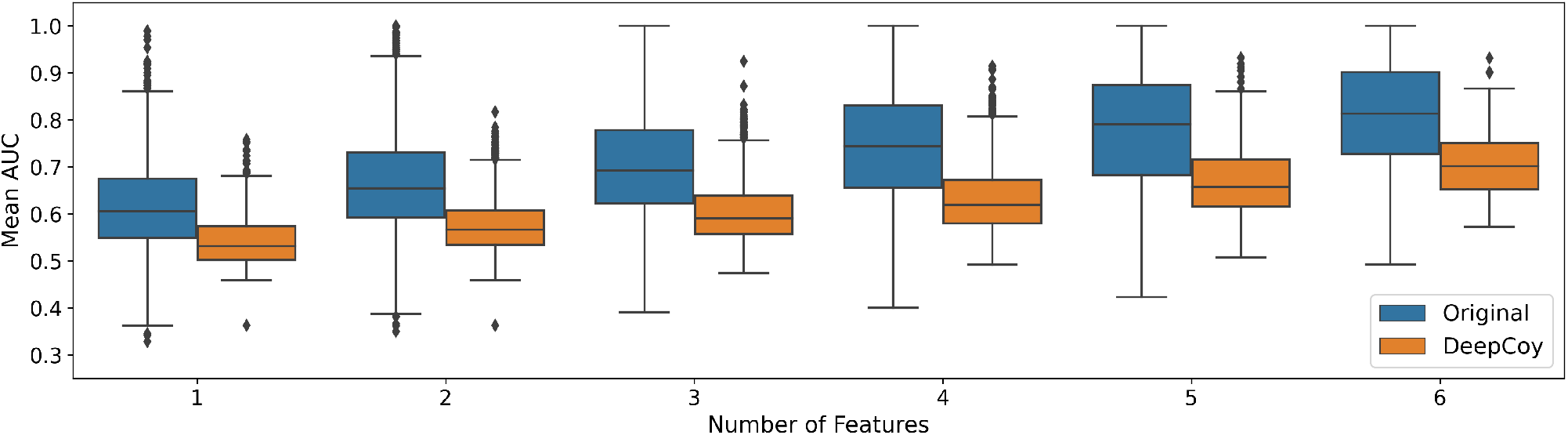
Results of the machine-learning based assessment of physicochemical property matching on DUD-E. Random forests were trained to predict whether a compound was an active or a decoy based on the unbiased features. Virtual screening performance was assessed by AUC ROC for the original DUD-E decoys and DeepCoy generated decoys. The DeepCoy generated decoys resulted in a reduction in the median per-target AUC ROC using all 6 features from 0.81 to 0.70 indicating a substantial reduction in bias.

However, even with the much improved property matching of the DeepCoy decoys, there remains some signal in the physicochemical properties. This is in part due to the high level of similarity between many of active molecules in DUD-E, a factor that should be controlled for when constructing the dataset to ensure low levels of bias (27). This is exemplified by the DUD-E target SAHH. DeepCoy decoys substantially reduced the DOE for the DUD-E properties to 0.11 (original decoys: 0.19). However, when assessing the decoys using the larger set of 27 properties, it became very challenging to unbias the decoy set (DeepCoy DOE: 0.29, original DOE: 0.34) due to high levels of similarity within the active set. All 63 active molecules for SAHH contain a similar fused ring system, while around half of the active molecules have 4 stereocenters (Figure 4). The considerable structural similarity, coupled with the high number of stereocenters for molecules of this size, was the primary cause of the poor DOE scores and is highly challenging to overcome via better decoy selection alone.

**Fig. 4.**
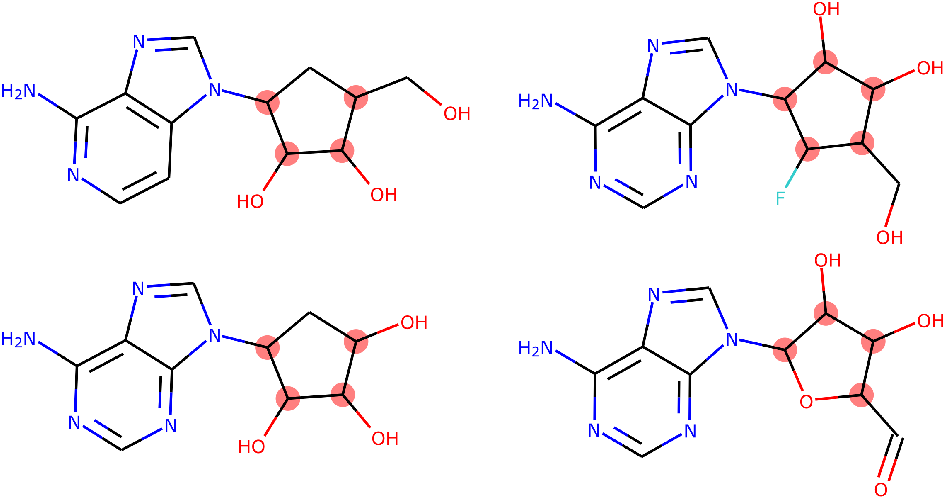
Four representative active ligands for DUD-E target SAHH. The 63 active molecules for SAHH have high levels of structural similarity, with all sharing similar fused rings systems. These ligands all have at least four stereocenters (highlighed in red, stereochemistry not shown), a property shared by over half of the active molecules for this target.

## 3.2. False Negative Bias

It is crucial that the improvement in property matching achieved by DeepCoy was not as a result of increasing the similarity between the active and decoy molecules, risking increasing false negative bias.

The average doppelganger score (26), a measure of the structural similarity between actives and decoys, remained consistent on the DUD-E set at 0.26 for the DeepCoy decoys and 0.25 for the original decoys, while the average maximum doppelganger score per target fell from 0.37 for the original decoys to 0.34 for the generated decoys. We saw similar results for the DEKOIS set; the average doppelganger score fell slightly (DeepCoy: 0.22, Original: 0.25), while there was a significant drop in maximum doppelganger score from 0.44 to 0.30 when using the DeepCoy decoys.

These results strongly suggest that the decoys generated by DeepCoy should not carry an increased risk of false negative bias compared to the original decoys.

### 3.3. Structure-based virtual screening

We further validated the quality of our generated decoys by docking the DUD-E set. Several publications have shown that most docking scoring functions are influenced by basic physicochemical properties (e.g. 2). In particular, Wallach and Lilien (28) showed that property mismatching can lead to an arbitrary increase *or* decrease in virtual screening performance of docking methods. Thus docking performance cannot be used alone to evaluate decoy molecules.

However, overall, better quality decoys should be harder to distinguish from active molecules, in particular if such decoys also more closely match the physicochemical properties of the active molecules and do not display an increased risk of false negatives.

The virtual screening performance of AutoDock Vina on the DUD-E set fell to an average per-target AUC ROC of 0.63 for the DeepCoy generated decoys compared to 0.71 for the original decoy molecules. There was a high correlation between the per-target docking performance using the original and DeepCoy decoys (Pearson’s R: 0.67, Figure S5) driven by the active molecules, which are common between both sets. However, for 83 of the 101 targets, the DeepCoy decoys led to a lower AUC ROC than the original decoys.

The decrease in the discriminative power of docking is likely driven by the closer property matching of the generated decoys, consistent with other studies (e.g. 26). For example, the original decoys for IGF1R resulted in a DOE score of 0.23, indicating a large mismatch between the active and decoy molecules. When this set was docked, Vina performed well with an AUC ROC of 0.81. In contrast, the DeepCoy generated decoys gave a DOE score of 0.02, a c. 90% reduction, and had a lower AUC ROC of 0.56. The inability for DeepCoy generated decoys to be easily separated from active molecules via docking together with the lack of additional risk of false negative is further validation of the suitability of these molecules for testing SBVS methods.

### 3.4. Synthethisability of generated decoys

A primary reason for selecting decoys from a virtual library of molecules is their high chance of synthetisability and the ability to purchase such compounds. However, for retrospective screening, or indeed training machine-learning models, decoys do not necessarily need to be synthetically feasible, but should be chemically possible (30).

A common criticism of molecules generated using *de novo* design methods is that they are not synthetically accessibile. We assessed the synthetic feasibility of molecules using the synthetic accessibility score (SA score, 4). The generated decoys have not been optimised for SA score nor selected based on this property. Despite this, the decoys generated by DeepCoy are, on average, relatively synthetically accessible, with an average SA score on the DEKOIS 2.0 set of 3.6 compared to 3.2 for the original decoys.

SA score is broadly a measure of molecular complexity, but with no regards to the precise functionality nor whether a given molecule should bind to a given target. Thus decoys should match the SA score (or a similar metric) of the active molecules, otherwise molecular complexity could become a distinguishing factor between actives and decoys.

As such, when generating decoys for DUD-E we included SA score as one of the properties to unbias. We demonstrate the effect this has on the SA score of decoy molecules by examining FA7, the median performing target (measured by DOE score) for the original decoy molecules, and NRAM, a target for which the active molecules have relatively high SA scores. The distributions of SA scores for FA7 and NRAM are shown in Figure 5 (mean SA score FA7 actives 2.9, NRAM actives 4.0). The DeepCoy decoys much more closely matched the SA score of the actives molecules of both targets than the original decoys, which did not match the SA score of the actives molecules in either case. This exemplifies the mismatch between SA scores of active and decoy molecules for some targets in DUD-E and demonstrates the adaptability of our generative framework.

**Fig. 5.**
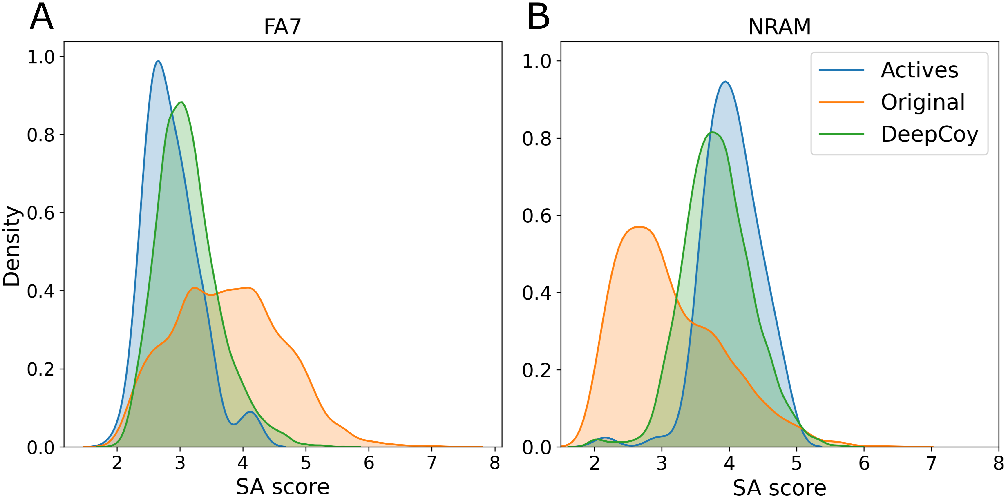
Synthetic accessibility (SA) scores for the active molecules (blue), original DUD-E decoys (orange), and DeepCoy generated decoys (green) for DUD-E targets FA7 (A) and NRAM (B). The DeepCoy generated decoys much more closely match SA scores of the active molecules than the original DUD-E decoys for both targets.

### 3.5. Effect of number of generated candidate decoys per active

We investigated how the number of candidate decoys generated per active with DeepCoy affects the quality of the final decoy set. Ideally as few candidates would be generated as possible; however, generating more candidates is likely to lead to a higher quality final decoy set. This creates a tradeoff between quality and computational requirements.

To explore this, we used the DEKOIS 2.0 target P38-alpha. This target achieved median performance as measured by DOE score with the original decoys, with a DOE score of 0.088 and doppelganger score of 0.22. We constructed multiple decoy sets by varying the number of candidate decoys generated by DeepCoy between 100 and 5000 per active molecule and selecting the best 30 as described previously.

Even generating only 100 candidates per active, the DOE score of the DeepCoy decoys was 0.079, representing an improvement over the original decoys of around 10%. As more candidates are generated, this difference rapidly increases (Figure 6), with a DOE score of 0.026 when 1000 candidates are generated, a 70% reduction compared to the original decoys. This continues to improve as more candidates are generated, albeit at a slower rate, reaching a score of 0.019 when 5000 are generated. The mean doppelganger score also decreased from 0.26 with 100 candidate per active to 0.23 when 5000 candidates were generated.

**Fig. 6.**
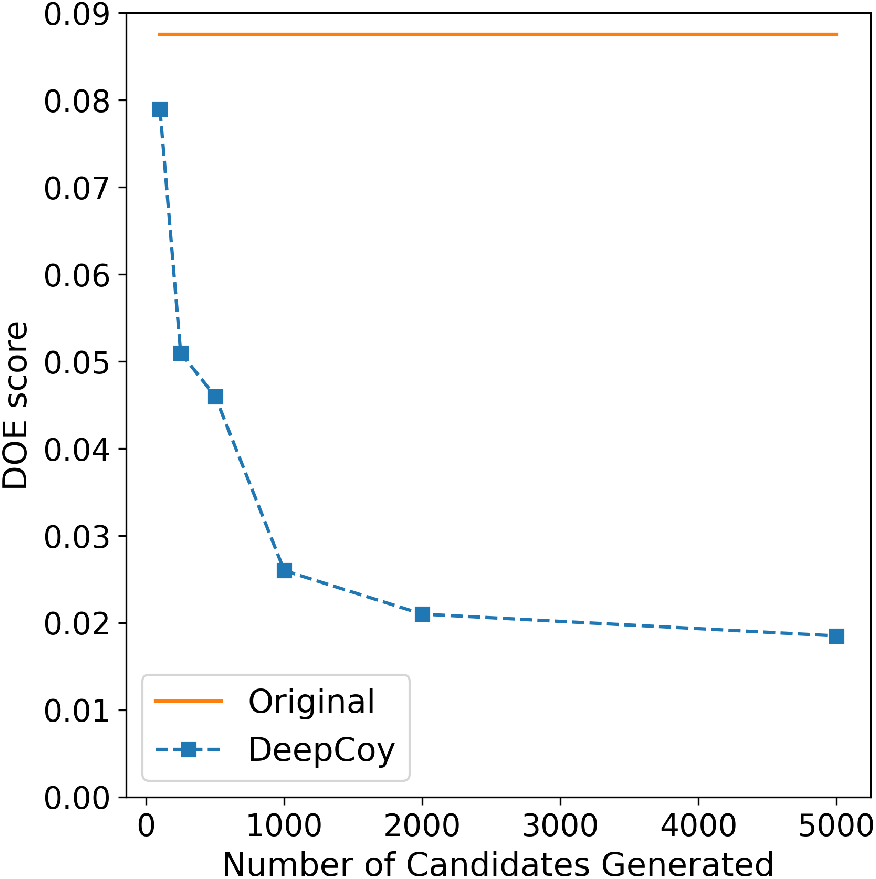
The effect on the DOE score of the final decoy set as the number of candidate decoys generated by DeepCoy is varied for DEKOIS target P38-alpha. In all cases, 30 decoys per active molecule are chosen. The DOE score for the DeepCoy generated decoys decreases rapidly as more candidates are generated, before slowing after 2000 potential decoys are generated. Even with only 100 candidates, the DOE score for the DeepCoy decoys is lower than the original decoys.

While there is a clear dependence between the quality of the final decoy set and the number of candidates generated, DeepCoy generated molecules outperformed the original decoys even when a very limited number of candidates were generated. Unlike a database approach where the maximum performance is limited by the dataset, in our framework the user can decide the desired level of property matching and risk of false negatives, generating additional candidate decoys until this is reached.

### 4. Conclusion

We have developed a graph-based deep learning method for generating property-matched decoy molecules for virtual screening. Unlike almost all virtual screening benchmarks, our method does not rely on a database molecules from which to select decoys but instead designs ones that are tailored to the active molecule.

We validated our generative model using two established structure-based virtual screening benchmarks, DUD-E and DEKOIS 2.0. For all 101 DUD-E targets and 80 of the 81 DEKOIS 2.0 targets, our generated decoy molecules more closely matched the physicochemical properties deemed by the respective datasets to be non-informative for binding, while introducing no additional false negative bias.

In particular, our generated decoys decreased the average DOE score from 0.163 to 0.032 for DUD-E and 0.109 to 0.038 for DEKOIS 2.0, an improvement of 81% and 66%, respectively. In addition, we demonstrated that they are no easier to distinguish than the original decoy molecules via docking with smina/Autodock Vina.

We believe that this substantial reduction in bias will benefit the development and improve generalisation of structure-based virtual screening methods. Currently, methods can perform well on retrospective benchmarks without performing molecular recognition by simply learning underlying biases (3, 22, 27). Thus it is unclear if improvements are genuine or due to more closely capturing these biases.

DeepCoy represents a novel approach to solve this problem, exhibiting substantial benefit over previous database-based methods. Our framework is highly customisable by the user and can naturally be combined with database search. While experimentally-verified inactives should be used whenever possible, this is not practically feasible apart from for limited-size benchmarking sets (e.g. 19). As such, effective decoys are crucial to the development of structurebased virtual screening methods.

The code is available at https://github.com/oxpig/DeepCoy. Generated molecules can be downloaded from http://opig.stats.ox.ac.uk/resources.

## Acknowledgements

The authors thank David Ryan Koes for providing docked protein-ligand poses of the original DUD-E dataset.

## References

1. Bauer, M. R. et al. (2013). Evaluation and optimization of virtual screening workflows with dekois 2.0 – a public library of challenging docking benchmark sets. J. Chem. Inf. Model., 53(6), 1447–1462.

2. Chaput, L. et al. (2016). Benchmark of four popular virtual screening programs: construction of the active/decoy dataset remains a major determinant of measured performance. J. Cheminf., 8(1), 56.

3. Chen, L. et al. (2019). Hidden bias in the dud-e dataset leads to misleading performance of deep learning in structure-based virtual screening. PLOS ONE, 14(8), 1–22.

4. Ertl, P. and Schuffenhauer, A. (2009). Estimation of synthetic accessibility score of drug-like molecules based on molecular complexity and fragment contributions. J. Cheminf., 1(1), 8.

5. Gómez-Bombarelli, R. et al. (2018). Automatic chemical design using a data-driven continuous representation of molecules. ACS Cent. Sci., 4(2), 268–276.

6. Huang, N. et al. (2006). Benchmarking sets for molecular docking. J. Med. Chem., 49(23), 6789–6801.

7. Imrie, F. et al. (2018). Protein family-specific models using deep neural networks and transfer learning improve virtual screening and highlight the need for more data. J. Chem. Inf. Model., 58(11), 2319–2330.

8. Imrie, F. et al. (2020). Deep generative models for 3d linker design. J. Chem. Inf. Model., 60(4), 1983–1995.

9. Jin, W. et al. (2019). Learning multimodal graph-to-graph translation for molecule optimization. International Conference on Learning Representations (ICLR).

10. Koes, D. R. et al. (2013). Lessons learned in empirical scoring with smina from the CSAR 2011 benchmarking exercise. J. Chem. Inf. Model., 53(8), 1893–1904.

11. Lagarde, N. et al. (2015). Benchmarking data sets for the evaluation of virtual ligand screening methods: Review and perspectives. J. Chem. Inf. Model., 55(7), 1297–1307.

12. Landrum, G. (2006). Rdkit: Open-source cheminformatics., [Online; accessed May 1, 2020).

13. Li, Y. et al. (2016). Gated Graph Sequence Neural Networks. International Conference on Learning Representations (ICLR).

14. Liu, Q. et al. (2018). Constrained Graph Variational Autoencoders for Molecule Design. Advances in Neural Information Processing Systems 31 (NeurIPS), pages 7795–7804.

15. Lyu, J. et al. (2019). Ultra-large library docking for discovering new chemotypes. Nature, 566(7743), 224–229.

16. Mysinger, M. M. et al. (2012). Directory of useful decoys, enhanced (DUD-E): Better ligands and decoys for better benchmarking. J. Med. Chem., 55(14), 6582–6594.

17. Polishchuk, P. G. et al. (2013). Estimation of the size of drug-like chemical space based on gdb-17 data. J. Comput.-Aided Mol. Des., 27(8), 675–679.

18. Rogers, D. and Hahn, M. (2010). Extended-connectivity fingerprints. J. Chem. Inf. Model., 50(5), 742–754.

19. Rohrer, S. G. and Baumann, K. (2009). Maximum unbiased validation (MUV) data sets for virtual screening based on PubChem bioactivity data. J. Chem. Inf. Model., 49(2), 169–184.

20. Réau, M. et al. (2018). Decoys selection in benchmarking datasets: Overview and perspectives. Front. Pharmacol., 9, 11.

21. Segler, M. H. S. et al. (2018). Generating focused molecule libraries for drug discovery with recurrent neural networks. ACS Cent. Sci., 4(1), 120–131.

22. Sieg, J. et al. (2019). In need of bias control: Evaluating chemical data for machine learning in structure-based virtual screening. J. Chem. Inf. Model., 59(3), 947–961.

23. Sterling, T. and Irwin, J. J. (2015). Zinc 15 – ligand discovery for everyone. J. Chem. Inf. Model., 55(11), 2324–2337.

24. Tran-Nguyen, V.-K. et al. (2020). Lit-pcba: An unbiased data set for machine learning and virtual screening. J. Chem. Inf. Model.

25. Trott, O. and Olson, A. (2010). AutoDock Vina: improving the speed and accuracy of docking with a new scoring function, efficient optimization and multithreading. J. Comput. Chem., 31(2), 455–461.

26. Vogel, S. M. et al. (2011). Dekois: Demanding evaluation kits for objective in silico screening — a versatile tool for benchmarking docking programs and scoring functions. J. Chem. Inf. Model., 51(10), 2650–2665.

27. Wallach, I. and Heifets, A. (2018). Most ligand-based classification benchmarks reward memorization rather than generalization. J. Chem. Inf. Model., 58(5), 916–932.

28. Wallach, I. and Lilien, R. (2011). Virtual decoy sets for molecular docking benchmarks. J. Chem. Inf. Model., 51(2), 196–202.

29. Wójcikowski, M. et al. (2017). Performance of machine-learning scoring functions in structure-based virtual screening. Sci. Rep., 7(March), 46710.

30. Yuriev, E. (2014). Challenges and advances in structure-based virtual screening. Future Medicinal Chemistry, 6(1), 5–7.

31. Zhavoronkov, A. et al. (2019). Deep learning enables rapid identification of potent ddr1 kinase inhibitors. Nat. Biotechnol., 37(9), 1038–1040.

