## Supplementary Material for "Generating Property-Matched Decoy Molecules Using Deep Learning"

### Additional DeepCoy model details

#### Atom types.

In line with both Liu et al.[1] and Imrie et al.[2], 14 atom types are permitted: carbon, nitrogen ($N^-$, N, $N^+$), oxygen ($O^-$, O, $O^+$), fluorine, chlorine, bromine, iodine, and sulphur (maximum valence 2, 4, or 6).

#### Network architecture.

Following Liu et al.[1], both the encoder and decoder utilise standard gated graph neural networks (GGNN),[3] which propagate messages for 7 steps and have residual connections between odd numbered time steps.

The neural network mapping the hidden state of a node to its atom type is implemented as a linear classifier, following Liu et al.[1].

#### Hyperparameters.

We trained the model with a learning rate of 0.001 for 10 epochs using the Adam optimiser and a batch size of 8, selecting the model from the epoch with the lowest validation loss. The dimension of the latent space was 100 and the dimension of the vector used to encode the target molecule during training and sampled from a $\mathcal{N}(\mathbf{0}, \boldsymbol{I})$ distribution when generating new molecules was 8. $\lambda_{KL}$ was set at 0.3.

We performed limited hyperparameter optimisation. However, we found that the model required a larger latent space dimension than in Imrie et al.[2], with the dimension of the latent space matching Liu et al.[1]. We believe this is due to the additional number of steps required to construct a full molecule compared to a partial structure as well as the importance of the input molecule for the generation task.

### Physicochemical properties to unbias

This section contains a list of properties that were unbiased. We calculated the values for all properties using RDKit.[4] This will likely cause minor discrepancies with the property values calculated in the construction of both DUD-E[5] and DEKOIS 2.0.[6] Indeed, values of the metrics used to assess property-matching and structural similarity were similar but not identical to the values reported by Bauer et al.[6]. All values reported here are calculated in the same way and thus directly comparable.

#### DUD-E.

DUD-E[5] selected the following six properties to match: molecular weight, log P, number of hydrogen bond acceptors, number of hydrogen bond donors, number of rotatable bonds, and net charge.

#### DEKOIS 2.0.

DEKOIS 2.0[6] was constructed by matching the following eight properties: molecular weight, log P, number of hydrogen bond acceptors, number of hydrogen bond donors, number of rotatable bonds, number of aromatic rings, positive charge, and negative charge.

#### All properties.

In our experiments using the DUD-E data set, we trained our model to match a substantially larger number of properties to assess whether DeepCoy could handle higher-dimensional restrictions.

Chaput et al.[7] focused their analysis of the DUD-E data set on nine properties, largely overlapping with the original DUD-E properties. In particular, they noted deviation between the DUD-E actives and decoys for the embranchment count and polar surface area.

We included these properties as the number of chiral centers (correlation of 0.97 with embranchment count) and topological polar surface area (TPSA),

MUV[8] was constructed to unbias the following 17 properties: simple counts of all atoms, heavy atoms, boron, bromine, carbon, chlorine, fluorine, iodine, nitrogen, oxygen, phosphorus, and sulfur atoms, number of hydrogen bond acceptors, number of hybond donors, logP, number of chiral centers, and number of rings.

Combining the properties used to construct DUD-E, DEKOIS 2.0, and MUV, together with two additional properties from Chaput et al.[7], synthetic accessibility,[9] and quantitive estimation of druglikeness (QED)[10] yields 27 unique phyiscochemcial properties.

### Additional results

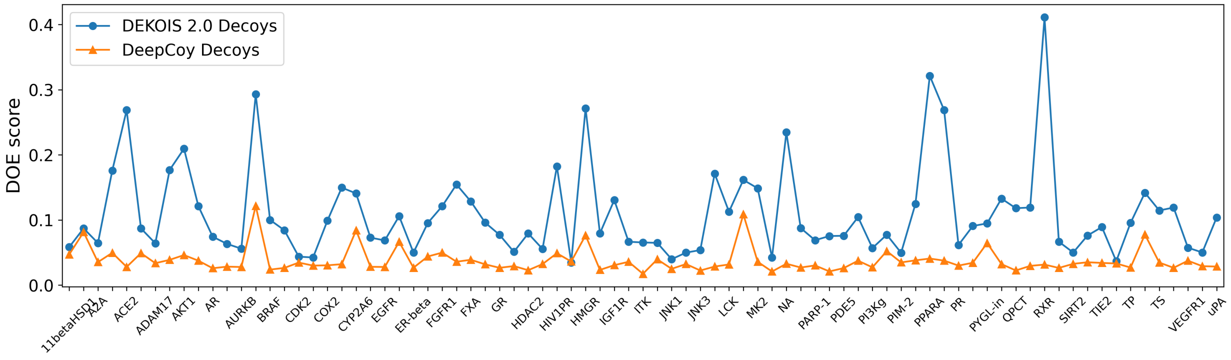


Figure S1: DOE scores of the original DEKOIS 2,0 set (blue) compared to the DeepCoy generated decoys (orange). The x-axis displays each DEKOIS 2.0 target in the same order as they appear in the DEKOIS 2.0 database (http://www.dekois.com/). The targets with even indices are not labeled on the x-axis due to space limitations.

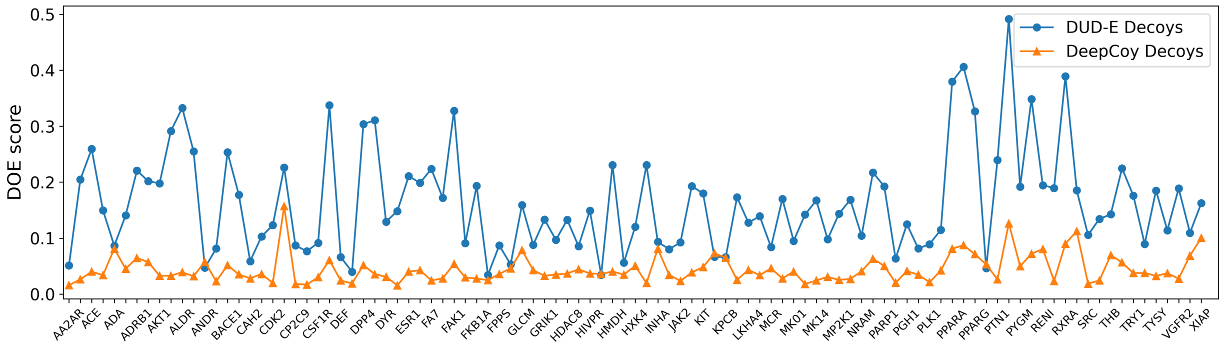


Figure S2: DOE scores of the original DUD-E set (blue) compared to the final DeepCoy generated decoys (orange) that were selected based on a larger number of properties to unbias, calculating DOE score using only the original DUD-E properties. The x-axis displays each DUD-E target in the same order as they appear in the DUD-E database (http://dude.docking.org/targets), excluding FPPS. The targets with even indices are not labeled on the x-axis due to space limitations.

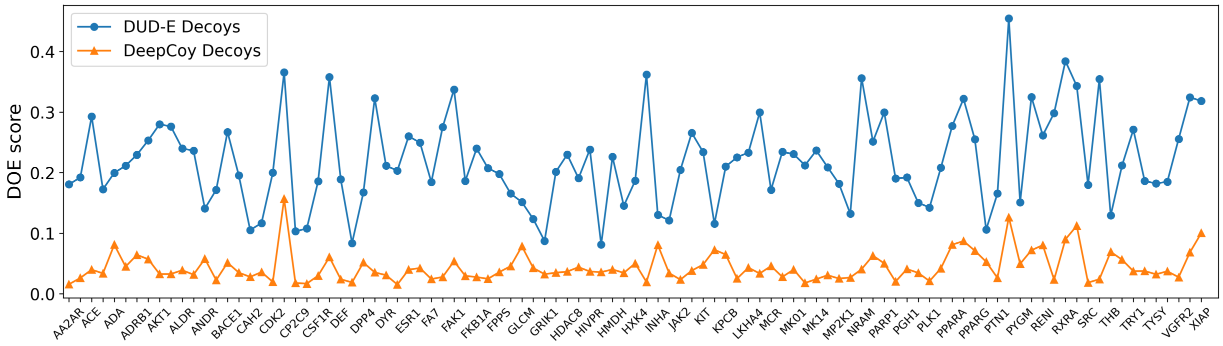


Figure S3: DOE scores of the original DUD-E set (blue) compared to the final DeepCoy generated decoys (orange) that were selected based on a larger number of properties to unbias, calculating DOE score using all 27 properties to unbias. The x-axis displays each DUD-E target in the same order as they appear in the DUD-E database (http://dude.docking.org/targets), excluding FPPS. The targets with even indices are not labeled on the x-axis due to space limitations.

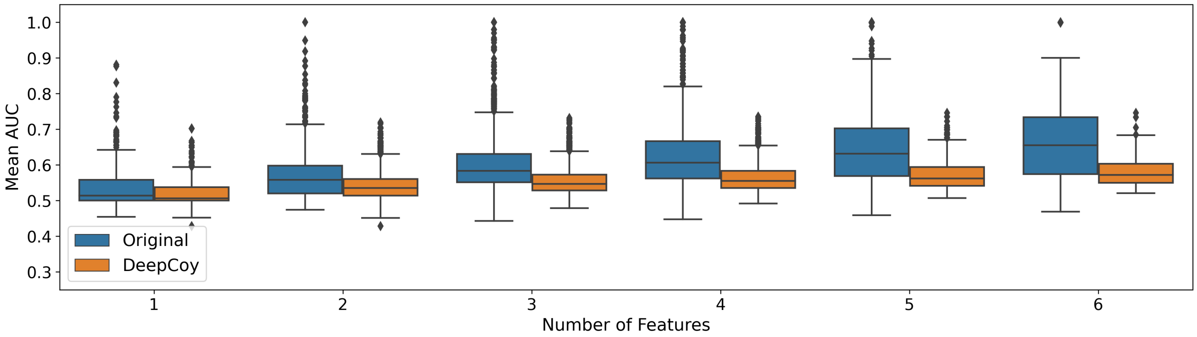


Figure S4: Results of the machine-learning based assessment of phyiscochemical property matching on DUD-E. 1-nearest neighbour models were trained to predict whether a compound was an active or a decoy based on the unbiased features. Virtual screening performance was assessed by AUC ROC for the original DUD-E decoys and DeepCoy generated decoys. The DeepCoy generated decoys resulted in a reduction in the median per-target AUC ROC using all 6 features from 0.66 to 0.57, indicating a substantial reduction in bias.

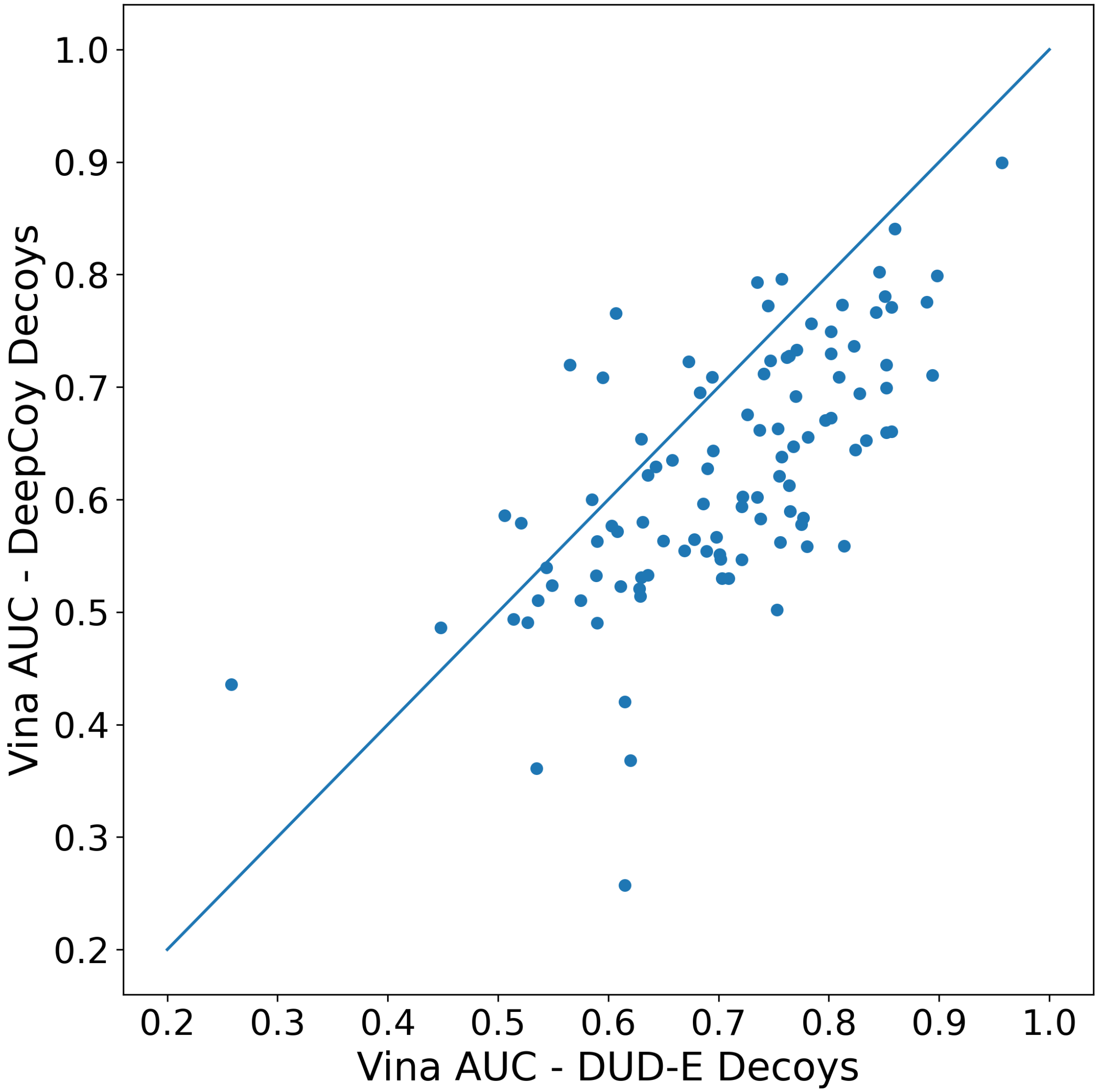


Figure S5: AUC ROC scores from docking with AutoDock Vina. The average AUC ROC decreased from 0.71 for the original DUD-E set to 0.63 for the DeepCoy generated decoys.

### References


(1) Liu, Q.; Allamanis, M.; Brockschmidt, M.; Gaunt, A. Constrained Graph Variational Autoencoders for Molecule Design. *Advances in Neural Information Processing Systems 31 (NeurIPS)* **2018**, 7795–7804.

(2) Imrie, F.; Bradley, A. R.; van der Schaar, M.; Deane, C. M. Deep Generative Models for 3D Linker Design. *J. Chem. Inf. Model.* **2020**, *60*, 1983–1995.

(3) Li, Y.; Tarlow, D.; Brockschmidt, M.; Zemel, R. Gated Graph Sequence Neural Networks. *International Conference on Learning Representations (ICLR)* **2016**,

(4) Landrum, G. RDKit: Open-Source Cheminformatics. , [Online; accessed May 1, 2020), 2006.

(5) Mysinger, M. M.; Carchia, M.; Irwin, J. J.; Shoichet, B. K. Directory of useful decoys, enhanced (DUD-E): Better ligands and decoys for better benchmarking. *J. Med. Chem.* **2012**, *55*, 6582–6594.

(6) Bauer, M. R.; Ibrahim, T. M.; Vogel, S. M.; Boeckler, F. M. Evaluation and Optimization of Virtual Screening Workflows with DEKOIS 2.0 – A Public Library of Challenging Docking Benchmark Sets. *J. Chem. Inf. Model.* **2013**, *53*, 1447–1462.

(7) Chaput, L.; Martinez-Sanz, J.; Saettel, N.; Mouawad, L. Benchmark of four popular virtual screening programs: construction of the active/decoy dataset remains a major determinant of measured performance. *J. Cheminf.* **2016**, *8*, 56.

(8) Rohrer, S. G.; Baumann, K. Maximum unbiased validation (MUV) data sets for virtual screening based on PubChem bioactivity data. *J. Chem. Inf. Model.* **2009**, *49*, 169–184.

(9) Ertl, P.; Schuffenhauer, A. Estimation of Synthetic Accessibility Score of Drug-Like Molecules Based on Molecular Complexity and Fragment Contributions. *J. Cheminf.* **2009**, *1*, 8.

(10) Bickerton, G. R.; Paolini, G. V.; Besnard, J.; Muresan, S.; Hopkins, A. L. Quantifying the chemical beauty of drugs. *Nat. Chem.* **2012**, *4*, 90–98.
